# Intravenous BCG immunization drives naïve CD4^+^ T cell to Th1 differentiation via CD4^+^ T cell epigenetic reprogramming

**DOI:** 10.1101/2025.05.23.655674

**Authors:** Ruilin Wang, Yuanhui Zhao, Jie Yang, Renjie Luo, Weikang Sun, Mengyu Zhang, Xiangdong Liu, Yayi Hou, Peng Cao, Erguang Li

## Abstract

Intravenous (IV) administration of live Bacille Calmette–Guérin (BCG) boosts potent immunity through coordinated interactions of innate immune response and adaptive immune response for BCG antiviral activity. The underlying mechanisms by which BCG initiates CD4^+^ T response remain incompletely understood. Here, we show that IV administration of BCG drives naïve CD4^+^ T cell differentiation to Th1 cell through epigenetic reprogramming. Single-cell transcriptomic and epigenomic analyses revealed that IV BCG immunization induced metabolic reprogramming in monocytes and increased the proportion of T cells. The immunization induced chromatin openness for transcription factors regulating naïve CD4^+^ T cells differentiation into Th1 cells. Importantly, we observed increases in Stat1, Irf1 and interferon-stimulated genes such as *Igtp* in Th1 cells, leading to upregulation of interferon related gene expression for antiviral response by BCG immunization. Activated transcription factors like Irf1 in CD4^+^ T cells promotes *Il12rb1* expression to facilitate IL-12 signaling and drive naïve CD4^+^ T cells differentiation into Th1 cells. We showed that BCG immunization reduced respiratory syncytial virus (RSV) infection. Thus, transcription regulation through epigenetic reprogramming plays a critical role in BCG-induced CD4^+^ T cell differentiation and to the broad antiviral activity.

**Significance:** Bacille Calmette–Guérin (BCG) vaccination confers nonspecific protection against heterologous pathogens. Recent work demonstrated that coordinated interactions of innate immune and persistent CD4^+^ T cell response plays a critical role in BCG induced antiviral activity. In our study, we investigated BCG immunization on epigenetic reprogramming of CD4^+^ T cells and found that BCG immunization elicited antiviral activity by provoking T cell proliferation. We observed chromatin openness favoring naïve T cell differentiation to Th1 response. Importantly, we observed STAT1 and interferon regulatory factor-1 (Irf1) activation. These findings provide mechanistic insights into BCG-induced Th1 response through epigenetic reprogramming and highlight the potential of BCG in offering a long-term protection against emerging and reemerging pathogens.

**Graphic abstract:** 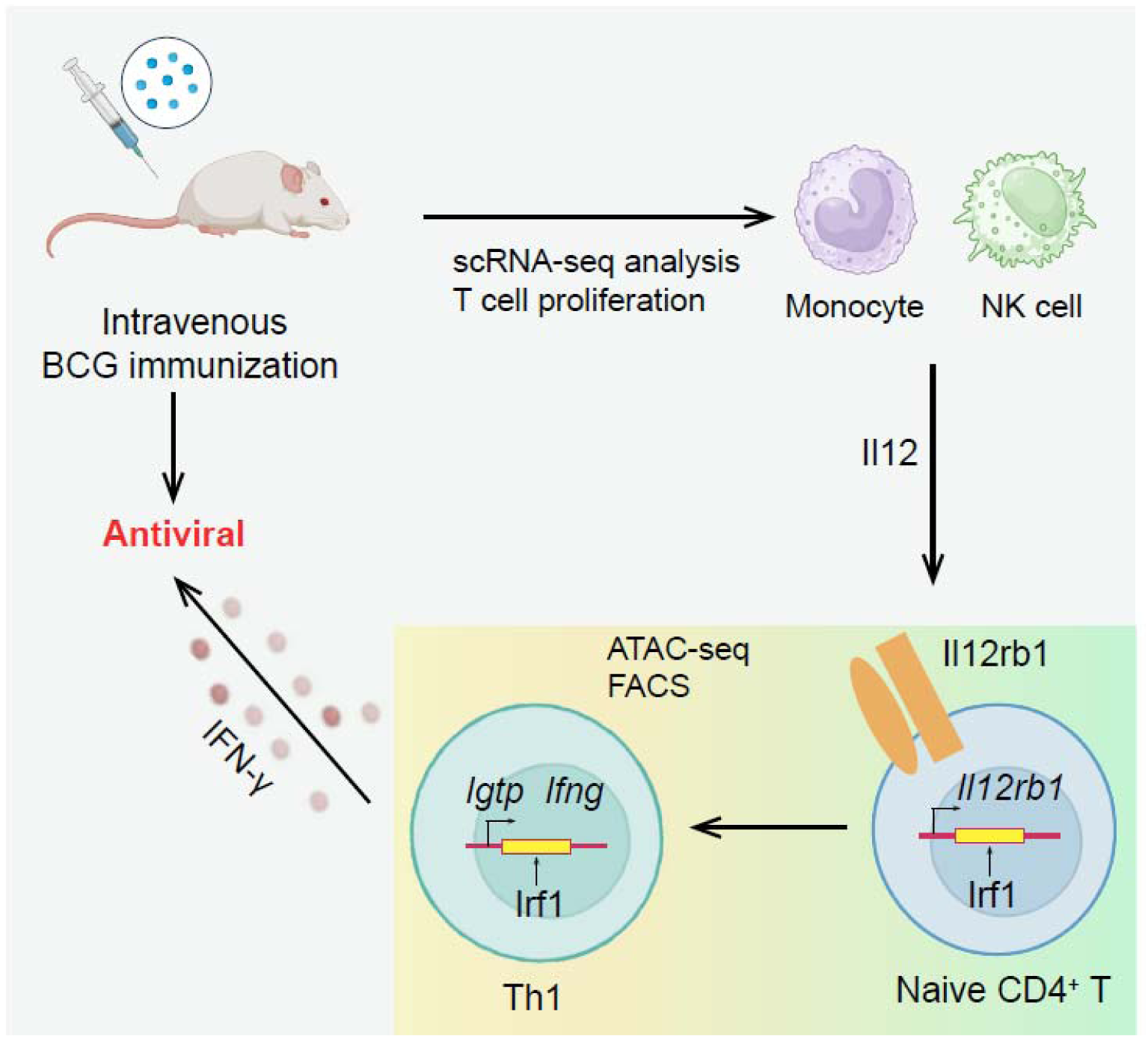

**Highlights:**

- Intravenous BCG immunization drives naïve CD4^+^ T cell differentiation to Th1 response;
- BCG alters chromatin accessibility for transcriptional factor expression favoring Th1 response through a Irf1-Il12rb1-IL12 feedback loop;
- The immunization elicits antiviral activity against RSV infection;

## Introduction

Bacille Calmette–Guérin (BCG) is a vaccine for tuberculosis discovered over a century ago and has been in clinical use ever since(1). In addition to its protective role against tuberculosis, BCG has been shown to confer broad nonspecific immunity, providing protection against various diseases. BCG vaccination by intradermal or intravenous (IV) provides protection against unrelated viral infections(2–4), including respiratory syncytial virus (RSV), influenza A virus, herpes simplex virus type 2 (HSV-2), and, recently, severe acute respiratory syndrome coronavirus 2 (SARS-CoV-2) by IV immunization(2). BCG vaccination also enhances host resistance to parasitic infections like malaria or schistosomiasis, as well as fungal and bacterial infections including *Candida albicans*, *Staphylococcus aureus*, and *Salmonella typhi* through T lymphocytes (T cells) independent immune responses(5–7). In addition, BCG vaccine has also been shown to confer beneficial effects against other diseases like cancer and remains as the most effective treatment for early-stage bladder cancer(8). Given its safety, stability, and broad protective effects, BCG holds significant potential for disease prevention and therapeutic applications. Elucidating a mechanism by which it provides broad protection against various diseases will provide valuable insights into immune-based disease interventions.

BCG is believed to exert its effects by activating the immune system to provide a broad-spectrum of antiviral effects(9). BCG vaccination induces protective integrated organ immunity through biphasic activation of innate and adaptive immune cells(10, 11). BCG vaccination promotes a robust antigen-specific type 1 helper T cell (Th1 cell) responses in the lungs(10). T cells orchestrate multiple aspects of adaptive immunity, including responses to pathogens, antigens, and tumors, making them indispensable for immune function. Therefore, depletion of CD4^+^ T cells prior to viral infection abolishes innate activation and protection by BCG vaccination(10). BCG vaccination alters the epigenetic landscape of progenitor cells in human bone marrow(12–14) and can induces the innate immune cells such as monocytes/macrophages and natural killer cells (NK cells) to undergo metabolic reprogramming through epigenetic modifications, thereby storing immune memory (trained immunity)(15, 16). Trained immunity bestows innate immune cells with non-specific protection, preventing re-infection, and is characterized by increased glycolysis and cytokine production in responsive cells(17, 18). It is therefore possible that the innate and adaptive immune interaction generates an enduring protective immunity against diverse pathogens. However, the immunological basis for this is unclear but T cell response has been speculated to play a major role in antiviral response(19).

In this study, we investigated IV BCG immunization on CD4^+^ T cells mediated immunity against RSV infection. We performed ATAC-seq on CD4^+^ T cells isolated from BCG-vaccinated mice and conducted a systematic analysis integrating publicly available single-cell datasets. Our findings reveal a pivotal role of epigenetic regulation in transcription factor (TFs) expression and in CD4 cell differentiation into Th1 cells for adaptive immune response. Our study provides epigenetic insights into BCG-induced epigenetic reprogramming in CD4^+^ T cell response.

## Results

### 1. Intravenous BCG immunization increases the proportion of T cells in the lung and spleen

It was recently reported that BCG vaccination stimulated integrated organ immunity by feedback of the adaptive immune response(10). To delineate a mechanism involved in adaptive cell response, we first referred to single-cell transcriptomic data (GSE244126) from the study(10). We integrated single-cell expression matrices from lung samples from BCG immunized at day 21 post BCG immunization and control groups (herein named D21 and control, respectively) and annotated cell types based on marker genes (Fig. 1A, 1B, and S1A). The proportion of T cells as well as NK cells and neutrophils was significantly increased at D21 compared to control (Fig. 1C). Cell-cell communication analysis revealed an overall enhancement in cellular interactions at D21 compared to control (Fig. 1D). Several immune-related signaling pathways were upregulated upon BCG vaccination (Fig. S1B). Notably, MHC class I and II signaling of T cell activation and function(20) was enhanced at D21. Additionally, we also observed upregulation of Bst2, an interferon-induced type II transmembrane protein that plays a key role in innate antiviral defense and T cell function(21, 22). These data suggested that BCG immunization induced immune cell proliferation and activation of antiviral gene expression.

**Figure 1.**
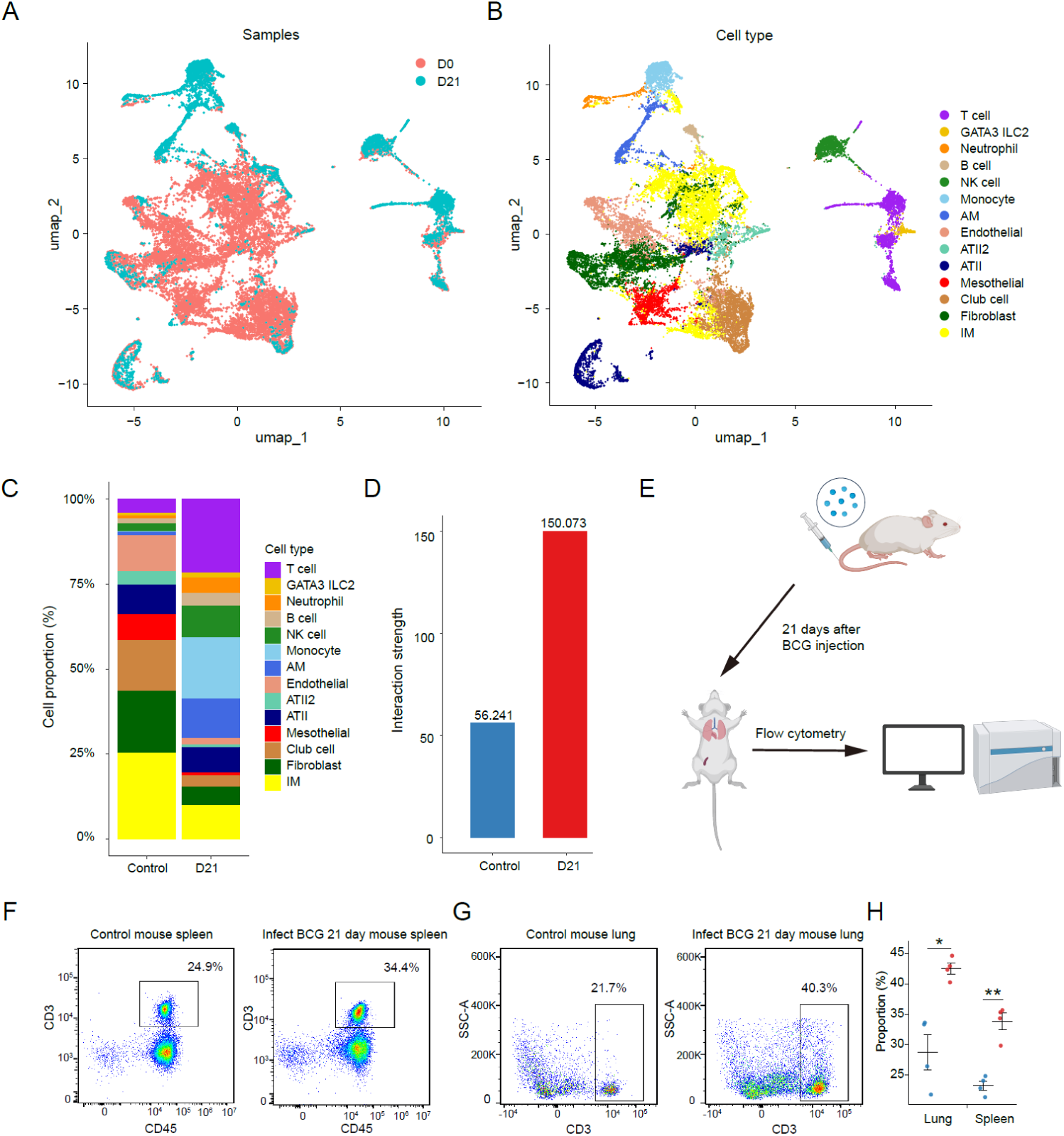
Single-cell transcriptomic data reveal an increased proportion of CD3⁺T cells in the lung and spleen at day 21 post BCG vaccination. **A**. UMAP (Uniform Manifold Approximation and Projection) plot showing the clustering results of GSE244126 samples, with different colors representing different BCG vaccination time points. **B.** Cell type annotation after clustering, with different colors representing distinct cell types. **C.** Bar plot showing the composition of different cell types at various time points post BCG vaccination, with different colors indicating different cell types. **D.** Bar plot representing the overall cell-cell interaction strength at different time points post BCG vaccination. **E-H.** Validation of BCG immunization on CD3^+^ T cell increase by flow cytometry. Balb/C mice were immunized by iv injection with 3×10^6^ CFU BCG (E). Control and immunized mice were killed and CD3⁺ T cells in the lung (F) and spleen (G) were analyzed (n=4) (H). * *p* < 0.05 and ** *p* < 0.01.

BCG immunization is known to drive monocytes to undergo metabolic reprogramming, a character of trained immunity(23). Specifically, GSEA pathway enrichment analysis revealed that the glycolysis pathway was enriched in monocytes at D21 post BCG immunization (Fig. S1C). Hexokinase (HK), a key enzyme regulating glycolysis(24), was notably upregulated in D21 monocytes (Fig. S1D), demonstrating IV BCG immunization boosted trained immunity.

We then performed experiments to validate these observations. In this regard, 6-week-old female Balb/c mice were injected with varying amount of BCG by intravenous (IV) route to induce immune response (Fig. 1E). Similar to those as reported, we detected increased cytokine production post BVG immunization (Fig. S1E and S1F). Flow cytometry analysis of the CD45^+^ cells in the spleen and lung samples collected at 21 days post BCG immunization showed that BCG immunization enhanced T cell population since higher proportion of CD3L T cells was detected in BCG immunized group (Fig. 1F-H and S1G).

### 2. Intravenous BCG immunization induces naïve CD4^+^ T cell differentiation into Th1 cells

To further explore the specific T cell subpopulations involved in antiviral immunity, we re-clustered T cells from the GSE244126 dataset based on T cell subtype marker genes, categorizing them into naïve T cells (*Sell*^+^, *Cd44*^−^), cytotoxic T cells (*Gzmb*^+^, *Prf1*^+^), Th1 cells (*Tbx21*^+^, *Infg*^+^), Th17 cells (*Rorc*^+^), and exhausted T cells (*Tox*^+^, *Tcf7*^+^) (Fig. 2A and S2A).

**Figure 2.**
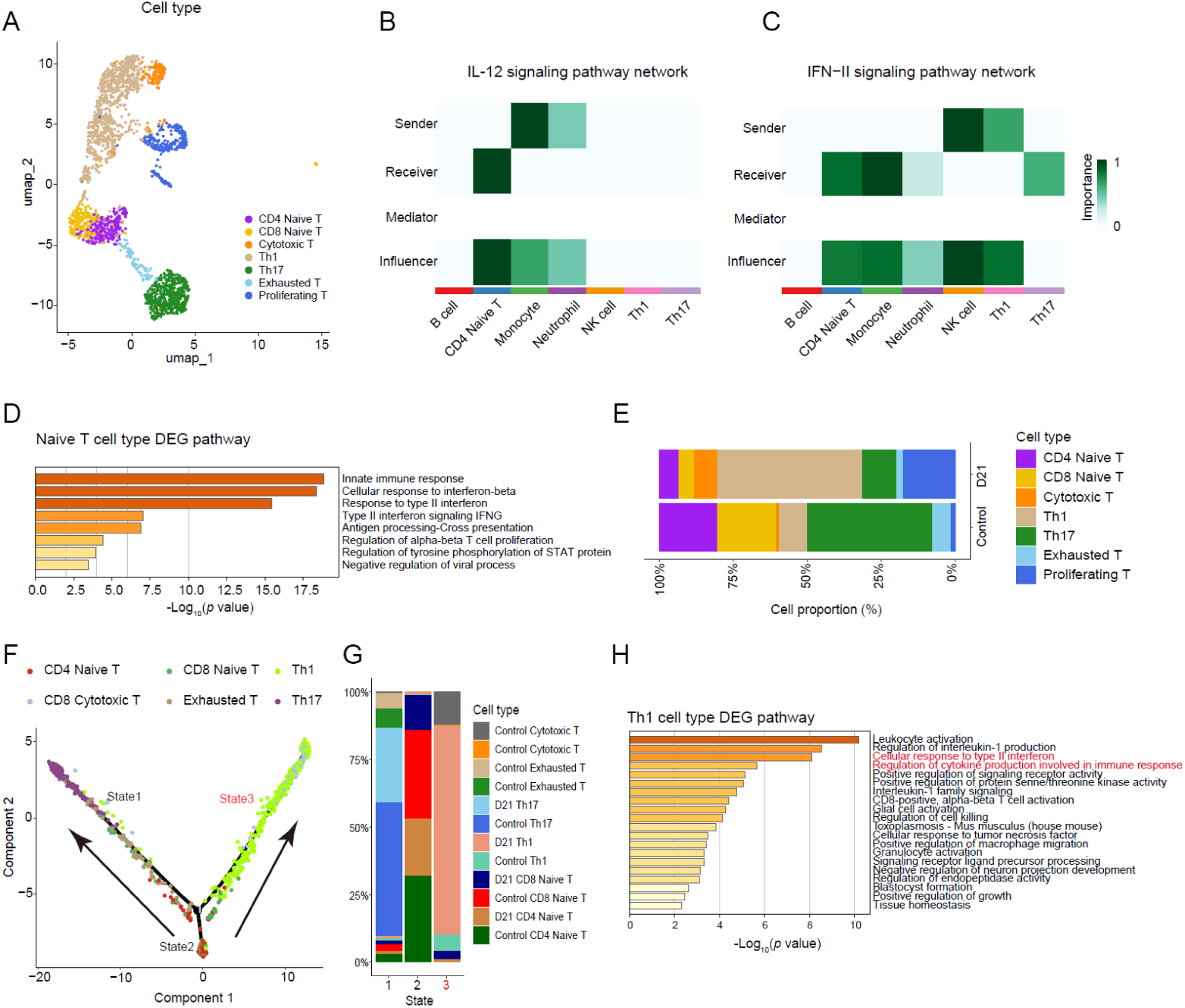
T cell re-clustering and differentiation of naïve CD4^+^ T cells into Th1 cells. **A.** The UMAP plot shows the re-clustering of T cells from the GSE244126 dataset, with cell types annotated based on T cell subtype marker genes. Different colors represent distinct T cell subtypes. **B-C.** IL-12 (B) and IFN-II (C) signaling pathway showing the centrality scores/importance of different cell types as senders, receivers, mediators, and influencers in the pathway. **D.** DEG enrichment analysis comparing naïve CD4^+^ T cells at D21 with control cells, highlighting pathways enriched with upregulated genes, with color representing *p*-value significance. **E.** Proportion of T cell subtypes at D21 and control, showing changes in T cell distribution across the two time points. **F.** Trajectory analysis of T cell differentiation, with arrows indicating the differentiation path of naïve T cells towards other subtypes. **G.** Bar plot showing the proportion of cell subtypes along the three differentiation trajectories of T cells. **H.** DEG enrichment analysis of Th1 cells, comparing D21 with control, focusing on pathways enriched in upregulated genes.

IL-12 signaling plays a crucial role in initiating T cell responses. Previous studies have shown that antigen-presenting cells release IL-12, activating T cells as one of the key factors in immune responses(25–28). Cell-cell communication analysis revealed that at D21, monocytes and neutrophils act as the sender cells in IL-12 signaling, while naïve CD4^+^ T cells are the receivers (Fig. 2B). Additionally, NK cells produce IFN-γ, which impacts naïve CD4^+^ T cell differentiation(29–31). In the IFN-II signaling network, Th1 and NK cells serve as the sender of signals, while monocytes and naïve CD4^+^ T cells act as receivers (Fig. 2C), indicating changes within CD4^+^ T cells after BCG immunization.

Differentially expressed gene (DEG) analysis of naïve CD4^+^ T cells at D21 compared to control showed enrichment in pathways related to “innate immune response”, “response to type II interferon”, and “regulation of tyrosine phosphorylation of STAT proteins” (Fig. 2D), which are associated with STAT family proteins and the role of IFN-II in stimulating naïve CD4^+^ T cell differentiation(32). Analysis of T cell subtype proportions revealed a decrease in naïve CD4^+^ T cells at D21 (Fig. 2E), a trend also observed in humans post BCG vaccination (Fig. S2B and S2C). In contrast, the proportion of proliferating T cells and Th1 cells increased (Fig. 2E), suggesting that naïve CD4^+^ T cells are activated and differentiated into Th1 cells, which play a significant role in antiviral responses.

Trajectory analysis revealed three stages of T cell differentiation (Fig. S2D). As time progresses, naïve T cells in stage 2 diverge into two branches, one of which is predominantly Th1 cells in BCG-treated mice at D21 (Fig. 2F and G, S2D). This further supports the idea that naïve CD4^+^ T cells differentiate into Th1 cells after BCG stimulation and contribute to antiviral immunity. Differentially expressed gene (DEG) enrichment analysis of Th1 cells showed upregulation of genes involved in leukocyte activation, regulation of cytokine production in immune responses, and cellular response to type II interferon (Fig. 2H).

These results indicate that naïve CD4^+^ T cells differentiate into Th1 cells upon receiving signals after BCG immunization, with Th1 cells likely playing a major role in BCG-induced innate immune memory and antiviral protection.

### 3. Intravenous BCG immunization on chromatin accessibility of CD4^+^ T cells

To further investigate these changes, we immunized mice with BCG by IV administration. The CD4^+^ T cells from the spleens of mock- or BCG-treated mice were isolated 21 days after immunization and used for ATAC-seq analyses (Fig. 3A, Fig. S3A). Flow cytometry analysis of IFN-γ in lung cells revealed an increased proportion of IFN-γ^+^ CD4^+^ cells in BCG-immunized mice and RSV-infected mice (Fig. S3B), consistent with the established role of Th1 cells in antiviral immunity through IFN-γ secretion. By enriching the TSS (transcription start site) regions, we observed good consistency across the samples, with consistent element enrichment across the samples, suggesting high data quality and minimal batch effects (Fig. S4C and S3D). Further correlation analysis of the read counts of the eight samples showed that the four BCG immunization samples clustered together, distinct from the control group, indicating group-specific differences between the two conditions (Fig. 3B). To compare the differential features between the groups, we performed differential peak analysis, which showed that each group had unique peaks and good consistency within each group (Fig. S3E). These differential peaks were then analyzed for element enrichment, revealing that BCG-immunized samples had upregulated peaks primarily in intron regions, whereas control samples had upregulated peaks in the distal intergenic regions (Fig. S3F). Next, we compared the gain and loss of peaks across the groups, and the results showed that the BCG group exhibited higher peak gain compared to the control group, suggesting a higher overall chromatin accessibility in CD4^+^ T cells after BCG immunization (Fig. 3C).

**Figure 3.**
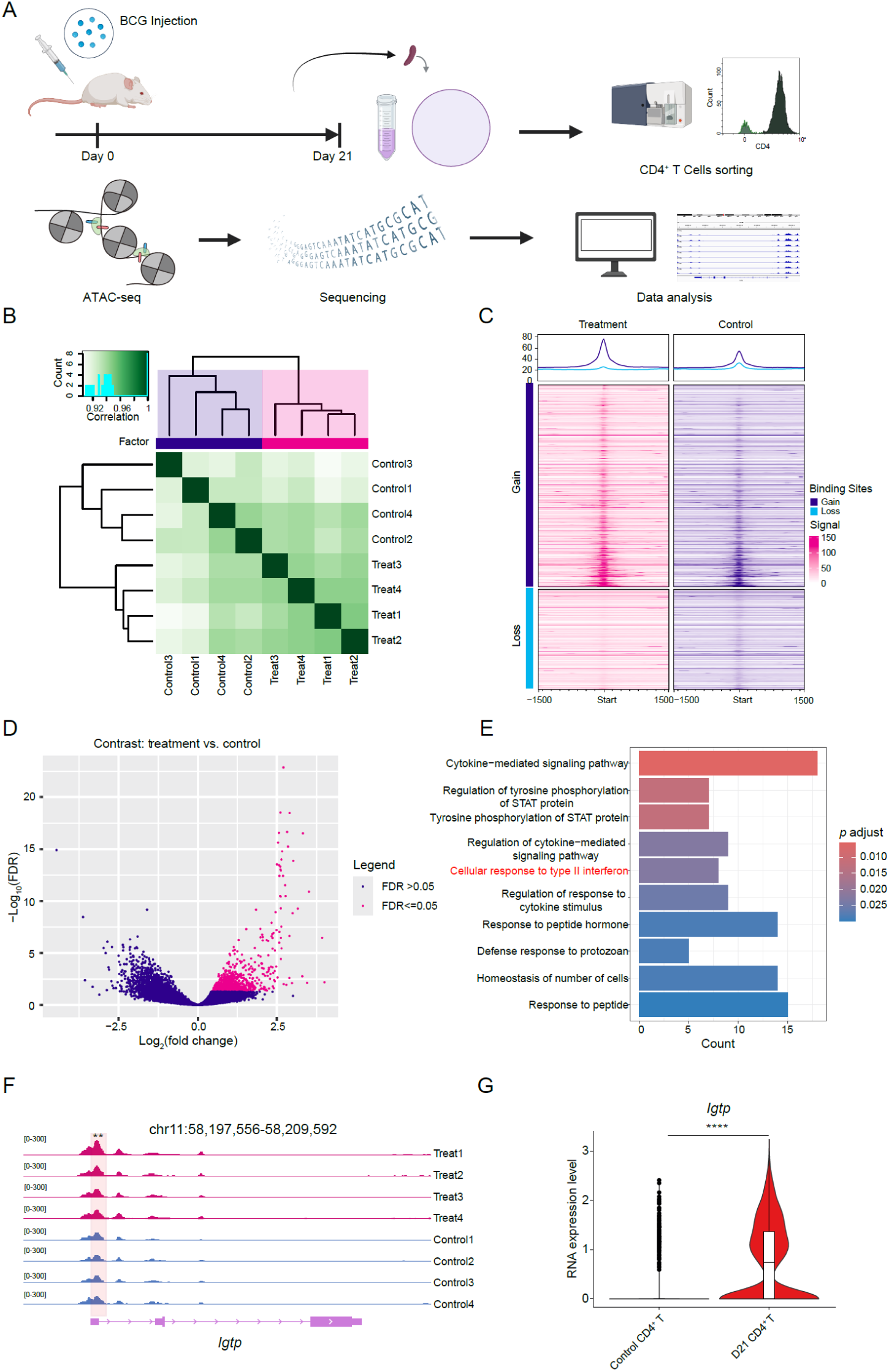
Chromatin changes in CD4^+^ T cells at D21 are associated with Th1 cell responses. **A.** Experimental workflow for ATAC-seq library preparation and sequencing, showing the steps involved in analyzing epigenetic changes in CD4^+^ T cells following BCG immunization. **B.** Heatmap of read count clustering for the BCG immunized samples (n=4) and control samples (n=4). The clustering shows distinct patterns between the two groups. **C.** Heatmap showing the gain and loss of peak regions in the two groups after sample merging. The signal strength differences indicate higher chromatin accessibility in the BCG group compared to the control group. **D.** Volcano plot illustrating the differential peaks between D21 and control. The pink markers indicate peaks that are upregulated in D21 and have a False Discovery Rate (FDR) ≤ 0.05, signifying their statistical significance. **E.** Pathway enrichment analysis of the pink peaks from the volcano plot, highlighting the “type II interferon signaling pathway” marked in red. This shows the key pathways activated in CD4^+^ T cells after BCG immunization. **F.** Chromatin accessibility of the *Igtp* across various samples. The promoter region with significant differences is highlighted in red circles. ** *p* < 0.01. **G.** Violin plot of *Igtp* expression levels in CD4⁺ T cells at the RNA level. **** *p* < 0.0001.

To further investigate chromatin changes in CD4^+^ T cells following BCG immunization, we focused on the significantly upregulated peaks from the BCG group compared to the control group (Fig. 3D). Upon annotation of these peaks to genes and performing pathway enrichment analysis, we found significant enrichment in pathways related to “STAT protein phosphorylation” and “cellular response to type II interferon” (Fig. 3E), consistent with our single-cell RNA-seq analysis (Fig. 2H), confirming that CD4^+^ T cells activated a Th1-mediated type II interferon response at both transcriptomic and chromatin accessibility levels.

By investigating the specific genes in these pathways, we identified interferon gamma induced GTPase (*Igtp*), a key IFN-γ regulatory gene(33), which appeared in both the cellular response to type II interferon and cytokine signaling pathways (Fig. 3E and S3G). The chromatin accessibility in the promoter region of *Igtp* was significantly elevated in CD4L T cells from the D21 group compared to the control group (Fig. 3F), and the expression of *Igtp* was significantly increased (Fig. 3G), indicating that *Igtp* was activated upon BCG immunization and contributed to the type II interferon response for IFN-γ production.

### 4. Intravenous BCG immunization activates Irf1 and regulates *Il12rb1* expression to promote naïve CD4^+^ T cell differentiation

Analyses of single-cell transcriptomic data uncovered TF changes among CD4^+^ T cell population. To further explore how these TFs function in the context of BCG-induced antiviral immune responses, enrichment of TF was predicted based on our ATAC-seq data. The results showed that interferon stimulated response element (ISRE) and interferon regulatory factor 1 (Irf1) exhibited increased enrichment in the D21 group compared to the control group, whereas enrichment of the transcription factor Ctcf was higher in the control group (Fig. 4A and 4B).

**Figure 4.**
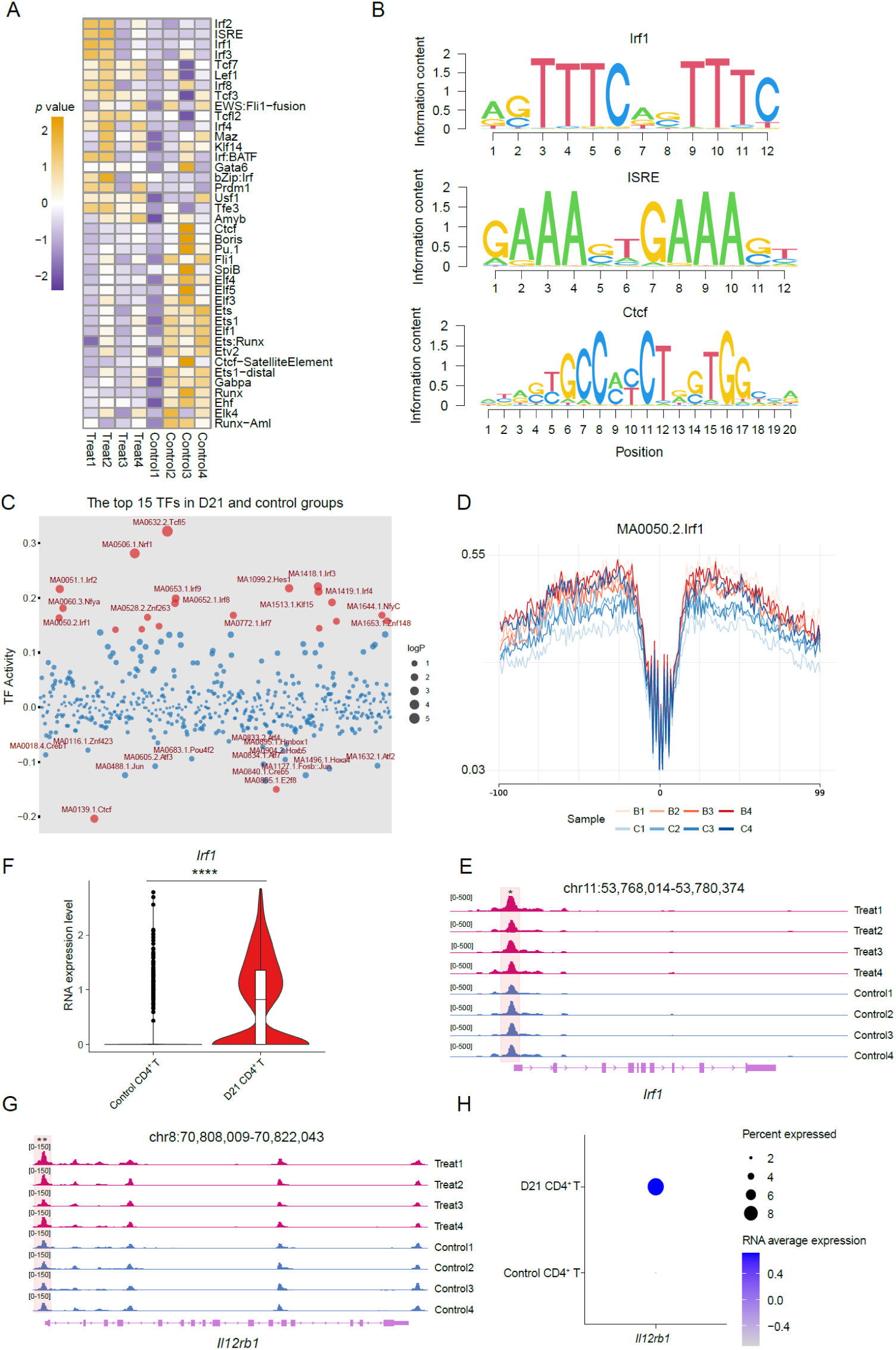
TF prediction and TF binding site footprint analysis. **A.** TF activity prediction in both groups. The heatmap displays the top TFs upregulated and downregulated in the D21 group compared to the control group. **B.** Motif analysis for TFs of Irf1, ISRE, and Ctcf in the D21 group, showing the binding motif. **C.** TF trajectory analysis for both groups. The top 15 TFs are sorted based on their TF activity. Circle size represents the *p*-value, with larger circles indicating smaller *p*-values, and red dots represent *p*-value< 0.05. **D.** TF activity of Irf1 in different samples, displayed to highlight its differential activity across conditions. **E and G.** Chromatin accessibility at *Irf1* (E) and *Il12rb1* (G), with significantly increased regions in the D21 group compared to the control group highlighted in red. * *p* < 0.05; ** *p* < 0.01. **F.** RNA expression levels of *Irf1* in CD4^+^ T cells, displayed as a vlnplot, showing the variation in expression across different samples. **** *p* < 0.0001. **H.** RNA expression levels of *Il12rb1* in CD4^+^ T cells, represented as a violin plot to compare the distribution of expression levels across the samples.

The transcription factor Irf1 is involved in many aspects of innate and adaptive immune responses by regulating target gene expression through interaction with its cognate recognition sequence ISRE in gene promoters. In the absence of Irf1, naïve CD4^+^ T cells fail to differentiate into IFN-γ-producing Th1 cells(34). We then performed TF footprint analysis on the combined samples, and the results showed that several IRF family members, including Irf1, were among the most active TFs in the D21 group (Fig. 4C, S4A). In each sample, Irf1 TF activity was higher in the D21 group than in the control group (Fig. 4D). In contrast, the top active TF in the control group was Ctcf (Fig. 4C), and its gene expression level was significantly lower in CD4^+^ T cells at D21 compared to the control (Fig. S4B). It has been reported that the complex formed by signal transducer and activator of transcription 1 (Stat1), Stat2, and Irf9 binds to ISRE, forming Isgf3, which serves as the primary driver of IFN-I-mediated transcription and is a crucial barrier to viral infections(35). Additionally, among the top 50 TFs with the largest variance in both groups, Stat1, Stat2, and Irf9 showed increased activity in the D21 group (Fig. S4C).

We then focused on the *Irf1*, which encodes the Irf1 protein. The chromatin accessibility in the *Irf1* promoter region was significantly higher in the D21 group compared to the control group (Fig. 4E), and the gene expression level was also higher in the D21 group (Fig. 4F). This suggests that *Irf1* plays a role in the immune response of CD4^+^ T cells following BCG immunization. Previous studies have shown that interferon induced Irf1 directly regulates the expression of *Il12rb1* and influences Th1 cell differentiation(36). In our analysis, we found that the chromatin accessibility in the *Il12rb1* promoter region was significantly higher in the D21 group compared to the control group (Fig. 4G). Moreover, the gene expression levels and the proportion of expression of *Il12rb1* were also higher in the D21 group (Fig. 4H). These results indicated that BCG immunization enhanced IL-12 signaling, which was consistent with our cell-cell communication analysis (Fig. 2B).

### 5. Intravenous BCG immunization increases regulatory activity of Stat1, Irf1 and interferon regulated genes in the Th1 cells

To further elucidate which CD4L T cell subsets are primarily responsible for the observed epigenetic changes, we carried out a gene expression regulatory network analysis using T cells from the GSE244126 dataset. The results showed an increase in the regulatory activity of Irf1 and Stat1 at D21 (Fig. 5A). Previous analyses indicated that Th1 cells play a crucial role in CD4^+^ T cell responses, so we specifically assessed the regulatory activity in Th1 cells. The results showed that these two regulatory factors (RFs) were among the top 5 regulators at D21 (Fig. 5B), and their gene expression levels were significantly upregulated in BCG treated mice compared to control (Fig. 5C), indicating their crucial role in the Th1 cell response induced by BCG.

**Figure 5.**
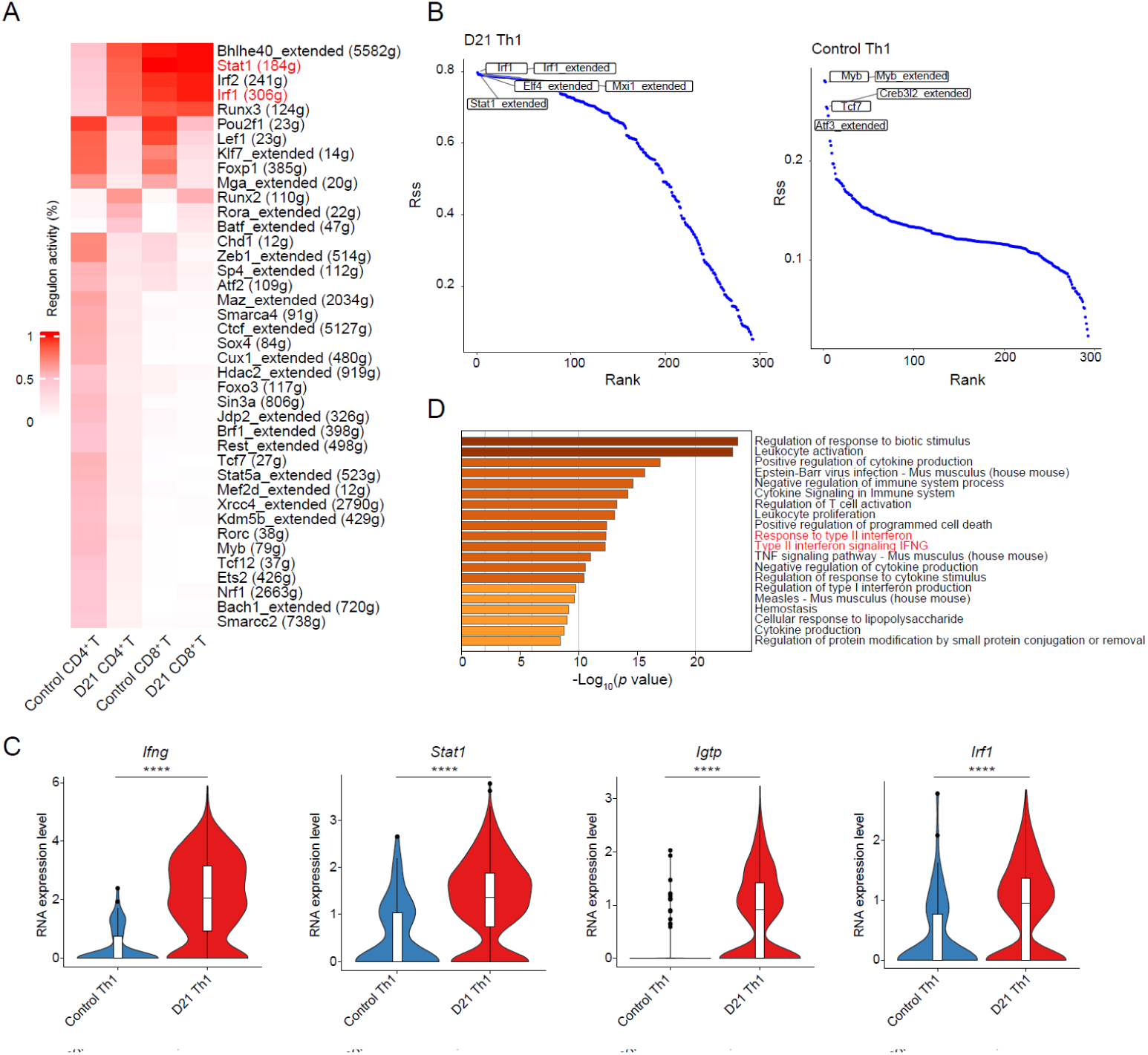
Gene expression network analysis of T cells reveals that Stat1 and Irf1 regulatory activity increases in Th1 cells. **A**. Heatmap of gene expression regulatory network activity for T cells, highlighting increased activity of Stat1 and Irf1 in CD4^+^ T cells at D21 compared to control. Red labels indicate upregulation of these RFs. **B.** Rssplot showing the regulatory specificity scores for Th1 cells at D21 and control after BCG immunization, with the top 5 RFs highlighted in the plot. **C.** Violin plot depicting the expression levels of genes enriched in the interferon related pathways, with statistical significance tested using the Wilcoxon test. The plot shows significant differences in gene expression between control and D21, with **** *p* < 0.0001. **D.** Pathway enrichment analysis of genes regulated by Irf1, with the IFN-II pathway prominently marked in red, indicating its activation.

Pathway enrichment analysis of genes regulated by Irf1 and Stat1 revealed enrichment in the “response to type II interferon” and “type II interferon signaling IFNG pathways” (Fig. 5D, Fig. S5A). In the Irf1 regulated genes, genes related to interferon responses such as *Ifng* and *Igtp* showed significantly higher expression levels at D21 to control (Fig. 5C). Similarly, proteins from the Gbp family, regulated by Stat1, including *Gbp3*, *Gbp4*, *Gbp5*, *Gbp6*, and *Gbp7*, were significantly upregulated in the type II interferon signaling pathway (Fig. S5B). In addition, *Irf1* expression was significantly elevated in naïve CD4L T cells at D21 compared to control (Fig. S5C). These changes in RNA expression likely reflect epigenetic alterations in CD4^+^ T cells, indicating that Irf1 not only mediates interferon responses by regulating genes such as *Igtp* in Th1 cells, but also plays a pivotal role in promoting the differentiation of naïve CD4L T cells towards the Th1 lineage. These findings highlight Irf1 and Stat1 in Th1 cells as key transcriptional regulators driving the type II interferon response following IV BCG administration, underscoring their essential role in the epigenetic reprogramming of CD4L T cells.

### 6. Intravenous BCG immunization protects mice against RSV infection

BCG administration provides superior protection compared to intramuscular and other conventional routes(10, 37). Finally, the effect of BCG on innate and adaptive immune activation was tested against RSV infection. To this end, BCG-immunized or mice in control groups were infected with RSV on the 21^st^ day post BCG immunization (Fig. 6A). Hematoxylin and eosin (H&E) staining of lung sections at 4 days post-infection revealed significant perivascular thickening in RSV-infected wild-type mice (RSV group). As a control, we also treated RSV-infected mice with ribavirin (RSV+Rib group), an approved antiviral drug for RSV treatment(38, 39). Notably, BCG-vaccinated mice exhibited reduced pulmonary viral loads (Fig. 6B-C), confirming the successful establishment of an RSV infection model. Consistent with recent reports with BCG immunization(10, 37, 40), mice receiving either BCG (BCG+RSV group) or ribavirin (RSV+Rib group) exhibited attenuated pathology, demonstrating that intravenous BCG administration boosted effective antiviral immunity against RSV infection.

**Figure 6.**
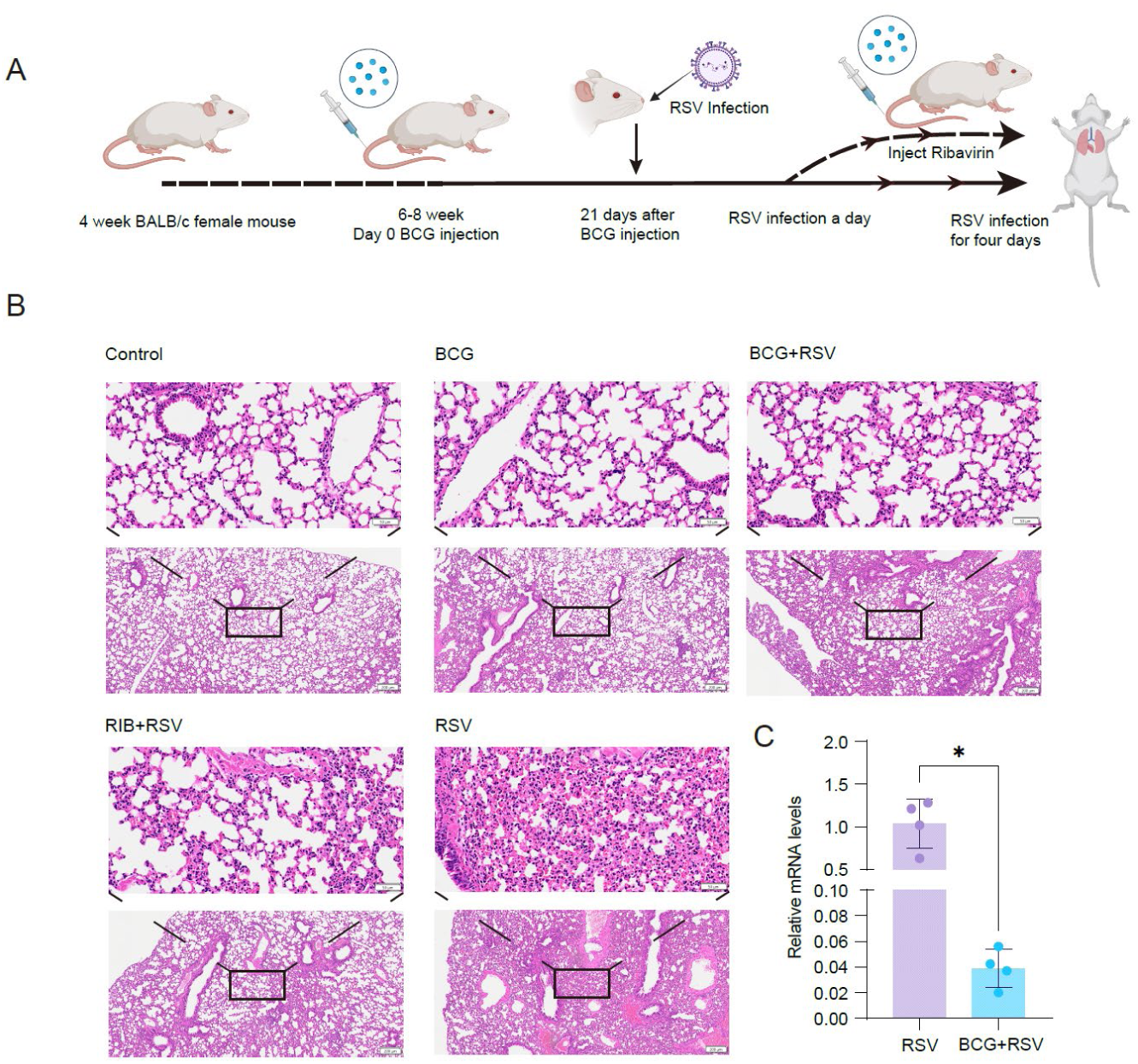
Intravenous BCG vaccination reduces RSV infection and lung injury in mice. **A.** Schematic representation of the experimental timeline for BCG vaccination and RSV infection in mice. **B.** H&E-stained lung sections from 5 experimental groups (Control, unimmunized and uninfected; BCG, BCG immunized not infected; RSV, RSV-infected; RSV+Rib, RSV-infected and Ribavirin-treated; BCG+RSV, BCG-immunized and RSV-infected). Black boxes indicate regions of interest that are magnified for detailed visualization. **C.** BCG immunization on viral load in the lungs. The RSV nucleoprotein gene (*N* gene) was reverse-transcribed and quantified by real-time qPCR. The data were normalized to GAPDH and are presented as mean ± SD (n=4). * *p* < 0.05.

In summary, data presented here demonstrated that BCG immunization enhanced CD4L T cell proliferation and transcription activation for naïve CD4L T cell differentiation and type II interferon expression against viral infection. This study thus demonstrates that IV BCG immunization provokes a Th1 response through epigenetic reprogramming favoring naïve CD4^+^ T cell differentiation to promote an interferon-mediated antiviral response.

## Discussion

BCG vaccination stimulates the adaptive immune response to imprint prolonged innate antiviral resistance against heterologous pathogens(1, 10). High-dose intravenous BCG vaccination induces enhanced immune signaling in the airways(41). A recent study showed that BCG IV elicited protection against SARS-CoV-2 and influenza 14 and 21 day post-vaccination that persisted for 3 months via both the innate and antigen-specific T cell responses following BCG immunization(10). In this study, we explored the role of epigenetic changes in CD4^+^ T cells in the antiviral effects of BCG. Through single-cell data analysis and validated by FACS method, we found that BCG administration enhanced the proportion of T cells in mice. Mechanistically, IV BCG induced a persistent transcriptional program connected to the stimulation of IL-12 and IFN-γ signaling and naïve CD4^+^ T cells differentiation into Th1 cells to combat against viral infections (Fig. 7).

**Figure 7.**
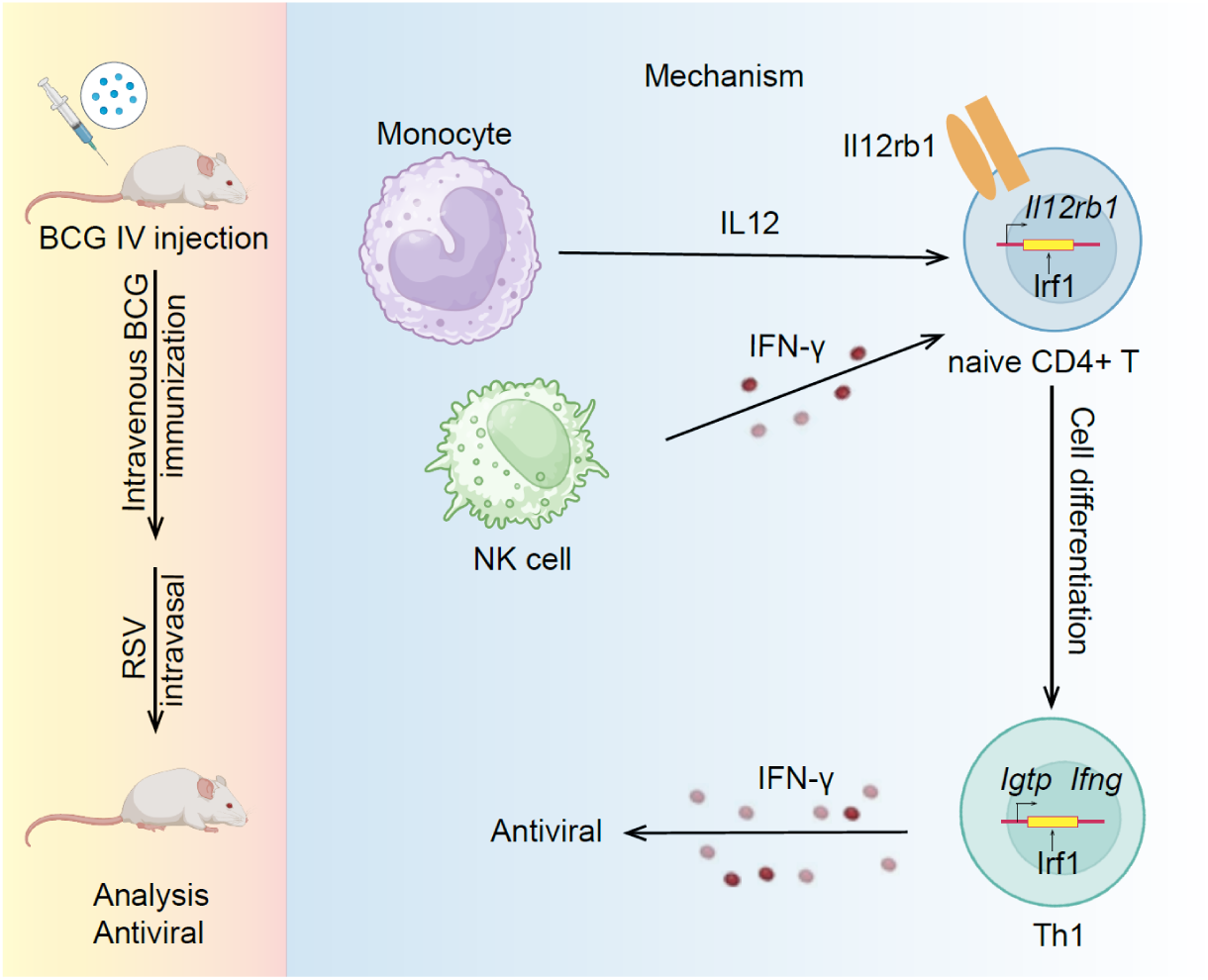
Possible mechanisms of BCG-induced transcriptional programming in CD4^+^ T cell differentiation and in protection against viral infections.

BCG is a Gram-positive bacterium. The host cells sense live mycobacteria and most purified mycobacterial antigens through Toll-like receptor 2 (TLR2). This pathway has been shown to be critical for the initiation of both innate and Th1 mediated responses(10). TLR2-deficient mice showed a 10-fold increase of pulmonary bacterial load 2 weeks post *Mycobacterium bovis* BCG infection, with a reduced adaptive response(42). TLRs have an important role in recognition of molecular signatures of invading microbe, leading to signaling pathway activation, and the adaptive immune response. TLRs utilize the extracellular leucine-rich repeat motifs to recognize pathogens while the cytoplasmic region, known as Toll/interleukin-1 receptor (TIR) domains, for heterophilic interactions with cytosolic adaptor proteins, such as MyD88 to initiate intracellular signaling events(43, 44). Thus, the protective effect of BCG against viral infections is attributed to MyD88-dependent signaling(10).

The Th1 cells secrete interferon (IFN) gamma and activates cytotoxic T lymphocytes (CTL) for clearance of viruses(45, 46). We performed ATAC-seq on CD4^+^ T cells from the spleens of mice administered BCG for 21 days, identifying global chromatin accessibility changes in CD4^+^ T cells after BCG treatment. Through combined single-cell transcriptomics and chromatin accessibility analysis, we highlight the important role of CD4^+^ T cells in this process. We identified Irf1, a key TF regulating Th1 cell differentiation, through combined analysis of TF footprints and gene expression regulatory network. Irf1 induces naïve CD4L T cells to differentiate into Th1 cells through the regulation of *Il12rb1* expression, promoting IFN-γ secretion for antiviral activity. Additionally, we observed enhanced CD4^+^ T cell proliferation along with upregulation of key regulatory genes like *Igtp*, which has been linked to Th1 cell-mediated IFN-II pathway and for immune response against viral infections(33).

Recent studies demonstrated that trained immunity does not work alone. Instead, it is regulated by T cell signaling(47), whereby antigen-stimulated CD4^+^ T cells feedback to imprint prolonged and broad innate-mediated antiviral resistance(10). The adaptive and innate immune systems cooperate within tissues to establish durable, antigen-independent protection against diverse pathogens(19). The monocytes from BCG immunized mice also exhibited metabolic reprogramming characterized by enhanced glycolysis and TCA cycle activity. Metabolic reprogramming is a hallmark of trained immunity(16). The broad, nonspecific protection induced by BCG has been attributed in part to metabolic and epigenomic remodeling in innate immune cells such as monocytes(48). Our data suggested that BCG induction of trained immunity is accompanied by a strong increase in glycolysis and, to a lesser extent, glutamine metabolism. We found that Irf7 and Stat1, critical regulators of both innate and adaptive immunity, exhibited increased transcriptional activity in monocytes. This led to enhanced IFN-α release and Hif-1α-mediated glycolytic responses, which influenced monocyte metabolic reprogramming and suggested that a shift of the glucose metabolism toward glycolysis is crucial for the induction of the histone modifications and functional changes underlying BCG-induced trained immunity against viral infection.

Our study contributes to the understanding of BCG actions through transcription reprogramming. BCG is typically given as an injection into the dermal tissue. The concept of immunization by alternative routes like aerosol or intravenous delivery of live attenuated *Mycobacterium bovis* to enhance protection against tuberculosis was suggested over 50 years ago. BCG immunization by IV administration is well-tolerated and effective in preventing tuberculosis infection in animals(49). Clinically, intravesical administration of BCG was one of the first FDA-approved immunotherapies and remains a standard treatment for bladder cancer. Thus, this study provides a mechanistic basis for the rational reapplication of BCG in clinical settings.

## Materials and Methods

### 1. Mouse model and housing conditions

BALB/c female mice (5 weeks old) were purchased from SPF Biotech (Beijing) and housed under specific pathogen-free (SPF) and temperature-controlled conditions. All animal experiments were conducted in accordance with ethical guidelines.

### 2. BCG immunization and RSV infection

BCG vaccine was obtained from Ruichu BIO (Lot: 20240718, Shanghai) and was resuspended in PBS prior to IV immunization. For the experiments, female BALB/c mice were immunized with 3x10^6^ CFU of BCG by IV route as previously reported(10). BCG-immunized or PBS-injected mice were randomly grouped and used on day 21 for RSV infection studies as we previously described(50), for immune cell analysis by flow cytometry, or for ATAC-seq library preparation of CD4^+^ T cells using a commercial kit (Vazyme, Nanjing). A list of key resources is provided as supplementary information, SI table 1.

### 3. Flow cytometry analysis of lung and spleen

Lung and spleen samples were collected from BCG-treated and wild-type BALB/c mice on day 21. Lung samples were enzymatically digested with 80 μl 100 μg/ml Liberase and 40 μl 0.2 mg/ml DnaseI for 40 min in 2 ml PBS at room temperature and mechanically dissociated through a 300-μm strainer to obtain single-cell suspensions. Place the spleen in a sieve and gently grind it with a tissue grinding rod. The mononuclear cells were subsequently purified by density gradient centrifugation after red blood cell lysis. Lung single-cell suspensions were stimulated with 10 ng/ml phorbol 12-myristate 13 acetate (PMA) and 1μg/ml Ionomycin in the presence of Brefeldin A for 6 h before membrane permeabilization. Lung and spleen cells were incubated with corresponding antibodies (APC-labeled anti-CD3, FITC-labeled anti-CD4, eFluor™ 450 labeled anti-CD45 and FIXABLE VIABILITY DYE EF780) at 4°C in the dark for 30 min with occasional mixing. Lung cells were harvested, fixed, permeabilized and stained with PE-Cy7-labeled anti-IFN-γ antibody at 4°C in the dark for 30 min. After washing with cold PBS, the cells were resuspended in Beckman Coulter CytoFLEX S flow cytometer and then analyzed on FlowJo (version 10.8.1) and CytExpert.

### 4. Histological analysis of lung tissues

Lung tissues were collected from four experimental groups: control, BCG-treated, RSV-infected (day 4 post-infection), and RSV-infected with ribavirin treatment. Samples were used for RNA extraction and RT-PCR analysis of viral gene expression or fixed in 10% neutral-buffered formalin. The tissues were embedded in paraffin, sectioned at 5 μm for hematoxylin and eosin (H&E) staining and pathological analysis.

### 5. Magnetic beads-based sorting of CD4**D** T cells

CD4L T cells were isolated from splenic single-cell suspensions using the Vazyme Mouse CD4L T Cell Isolation Kit (CS102, version 24.1). Briefly, spleens were mechanically dissociated using a 70-μm cell strainer, and red blood cells were lysed with 3 ml of lysis buffer for 10 min at room temperature. Cells were washed, resuspended in selection buffer (1 mM EDTA and 2% FBS), and passed through a 70-μm filter. The final cell concentration was adjusted to 1×10L cells/ml, and CD4L T cells were isolated using a magnetic separation rack.

### 6. ATAC-seq library preparation from splenic CD4**D** T cells

Purified CD4L T cells were used for ATAC-seq library construction using the Vazyme Hyperactive ATAC-Seq Library Prep Kit for Illumina (TD711, version 23.2), following the manufacturer’s protocol. Briefly, 5×10L cells/ml were lysed in Lysis Buffer, followed by transposition with 50 μl of fragmentation mix at 37°C for 30 min. DNA fragments were purified using ATAC DNA Extract Beads, and libraries were amplified for 13 cycles. The final libraries were purified using ATAC DNA Clean Beads before sequencing.

### 7. ATAC-seq data analysis

#### Preprocessing of raw data

Raw sequencing reads in FASTQ format were subjected to quality control using fastp software (version 0.24.0)(51). Reads were aligned to the mm10 reference genome (https://hgdownload.soe.ucsc.edu/goldenPath/mm10) using BWA software, followed by duplicate removal with Sambamba (version 1.0.1) software(52). The aligned reads were sorted using SAMtools(53), and bigWig format files were generated with BAMscale software for visualization in Integrative Genomics Viewer (IGV).

#### Identification of accessible regions and TF enrichment analysis

Peak calling was performed using the “callpeak” function in MACS2 software (version 2.2.6)(54) with the following parameters: not considering local bias at peak candidate regions and bypassing the shifting model. The sequencing fragment length was manually set to 75, with an extension length of 150. A *p*-value threshold of 0.05 was used to define statistically significant peaks.

For TF enrichment analysis, the HOMER software(55) was employed. The configureHomer.pl script was used to establish the mm10 HOMER database, and findMotifsGenome.pl was applied for motif discovery, with a region size set to 500 bp.

#### ATAC-seq transcription factor footprinting analysis

To identify transcription factor (TF) binding activity from bulk ATAC-seq data (GSE296985), we performed footprinting analysis using the HINT-ATAC algorithm(56). Tn5 insertion sites were inferred by adjusting read alignment positions to reflect transposase cutting sites. Peak regions were first identified using MACS2 with default settings and used as input for footprinting. Differential footprinting analysis between experimental groups was conducted using the “rgt-hint differential” module.

#### Visualization of TSS accessibility

BigWig files were generated using the bamCoverage function from DeepTools software (version 3.5.6)(57), with RPKM normalization to account for sequencing depth differences across samples. In R, the GenomicFeatures package was used to process the mm10 genome annotation file obtained from the UCSC genome browser. The TSS4K regions, were established using the “resize” function. To analyze signal distribution, the “computeMatrix” function from DeepTools software was employed in scale-regions mode to normalize all genomic regions to the same length. A bin size of 10 bp was specified for coverage calculation, followed by heatmap visualization to illustrate signal intensity across TSS regions.

#### Differential peak identification, annotation, and enrichment analysis

Differential peaks were identified using DiffBind (v.3.14.0) software (https://github.com/nshanian/DiffBind) across the different samples. The control and BCG-treated samples were grouped into two separate conditions for comparison. The “dba.contrast” function was used to define the contrast, followed by differential analysis using “dba.analyze” function, and the results were obtained using “dba.report” function with the EdgeR method. Differential peaks were annotated to the mouse mm10 reference genome using ChIPseeker software (v.1.40.0)(58). For the annotated genes, Gene Ontology (GO) enrichment analysis was performed using the clusterProfiler software (v.4.12.6)(59), with results sorted by *p*-value in ascending order. The top 10 enriched pathways were selected for visualization.

### 8. Single-cell transcriptomics

#### Cell clustering, identification, and differential gene pathway enrichment analysis

For the GSE244126 dataset, we performed clustering analysis using the Seurat package (version 5.1.0)(60). Following data import, we created a Seurat object and applied quality control filters based on published criteria: cells with RNA counts between 500 and 50,000 and mitochondrial content below 10% were retained. Data normalization was performed, and the top 2,000 highly variable genes were identified. Principal component analysis (PCA) was conducted, with the first 13 principal components selected for downstream analysis. We then constructed a shared nearest-neighbor graph and applied Louvain clustering with a resolution of 0.8, identifying 29 distinct clusters. Dimensionality reduction was performed using the “RunUMAP” function.

Cell-type annotation was conducted based on canonical marker genes. Following annotation, differentially expressed genes (DEGs) across cell types were identified using the “FindMarkers” function, with filtering criteria set as min.pct = 0.1, min.diff.pct = 0.1, and only.pos = TRUE. Identified DEGs were subsequently subjected to pathway enrichment analysis using the Metascape platform(61).

#### Cell-cell interaction analysis

Cell-cell communication was analyzed using the CellChat package (version 1.6.1)(62). Seurat objects corresponding to BCG vaccination at day 21 and control were used as inputs, with the “CellChatDB.mouse” database selected. The “mergeCellChat” function was used to integrate the two datasets. Interaction strength between control and D21 was assessed using the “compareInteractions” function with measure = "weight", while signaling pathway activity was compared using the “rankNet” function. Interaction networks were visualized using the “netVisual_bubble” function.

#### GSEA pathway enrichment analysis and GSVA score calculation

Gene set enrichment analysis (GSEA) was performed using the msigdbr package (version 7.5.1) to retrieve hallmark gene sets. Single-sample gene set enrichment analysis (ssGSEA) scores were calculated using the “ssgseaParam” function from the GSVA package(63), with T-cell expression data and hallmark gene sets as inputs. The “gsva” function was then applied to compute GSVA scores, which were visualized using the “pheatmap” function to generate heatmaps of GSVA scores across different T-cell subsets.

#### Gene regulatory network analysis

Gene regulatory network analysis was conducted using the SCENIC package (version 1.3.1)(64). First, putative transcriptional regulators were identified based on co-expression patterns using GENIE3 (version 1.12.0). Candidate target genes directly interacting with regulatory motifs were then identified using RcisTarget (version 1.10.0). Finally, the AUCell package (version 1.12.0) was used to assess the activity of gene regulatory networks at the single cell level.

### 9. Statistical Analysis

For DEG analyses of single-cell RNA-seq data, *p*-values were calculated by the Wilcoxon rank sum test. *, *p*-values <0.05; **, *p*-values <0.01; ***, *p*-values <0.001; ****, *p*-values <0.0001.

## Supporting information

By Wang, et al, Intravenous BCG immunization drives naive CD4+ T cell to Th1 differentiation via CD4+ T cell epigenetic reprogramming

## Acknowledgement

The study was supported by a grant from National Key R&D Program of China (2023YFC2308200, to Peng Cao, Yayi Hou and Erguang Li). We thank the authors for providing publicly accessible GSE244126 and PRJNA1095450 dataset.

## Authorship

RW and EL designed the study, interpreted the data and wrote the manuscript. RW and EL planned the research, RW and YZ analyzed the data. RW, RL and YZ performed immunization and infection experiments including challenge, viral load quantification and sample analysis. RW, YJ and MZ performed cell isolation and conducted ATAC library preparation. RW performed scRNA-seq and ATAC-seq analysis. RW and YZ immunized mice and performed flow cytometry analysis. RL and WS propagated the RSV virus and performed infection assay. XL and YH provided reagents. PC for funding acquisition. EL for supervision and administration. RW and EL wrote the paper. All authors have reviewed and edited the manuscript.

## Data availability statement

Source data are provided with this paper. The ATAC-seq data generated in this study were submitted to the GEO repository with an accession number of GSE296985 and will be publicly accessible upon acceptance of the paper. The scRNA-seq data of BCG-immunized mice (GSE244126) and CD3 positive populations from human PBMCs (PRJNA1095450) were retrieved from the NCBI.

## Code availability

Computer code is available upon reasonable request.

## Ethics statement

A protocol (D2312001) for the care and use of animals was reviewed and approved by IACUC committee of Nanjing University.

