## Supplementary material for "Intravenous BCG immunization drives naïve CD4^+^ T cell to Th1 differentiation via CD4^+^ T cell epigenetic reprogramming": By Wang, et al, Intravenous BCG immunization drives naive CD4+ T cell to Th1 differentiation via CD4+ T cell epigenetic reprogramming

Figure S1

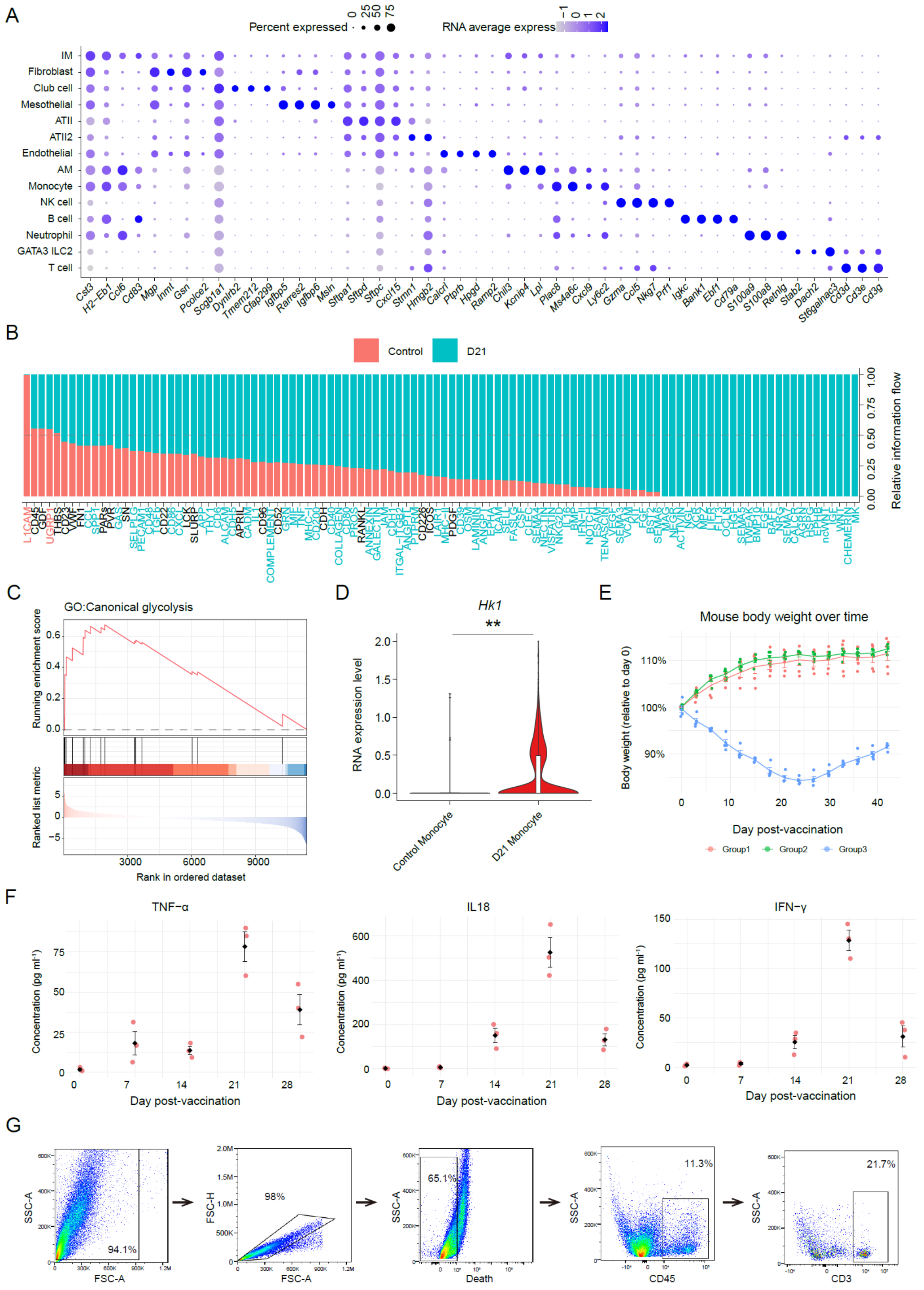


**Figure S1. Markers for cell type identification and flow cytometry gating strategy.**

**A.** Dot plot showing the expression of marker genes for different cell types, with color

intensity representing the average expression level of each gene, and circle size indicating the proportion of cells expressing the gene.
**B.** Comparison of cell signaling pathway strength between D21 (blue) and control (red) pathways. The analysis was performed using paired rank sum tests ($p < 0.05$).
**C.** GO enrichment analysis for the glycolysis pathway using GSEA.
**D.** The violin plot of RNA gene expression levels of *Hk1*. ** $p < 0.01$.
**E.** Determination of BCG dose for intravenous BCG immunization. Female BALB/c mice were immunized with BCG by IV immunization using 5 mice per dose group (red, $1x10^6$; green, $3x10^6$; blue $6x10^6$ CFU/mouse). Body weight was monitored on time as indicated.
**F.** Determination of cytokine production in BCG or mock immunized mice. The blood samples (n=3) were collected by retro-orbital bleeding from $3\times10^6$ CFU BCG immunized group and used for IL18, IFN-γ and TNF-α detection by ELISA (CUSABIO, Wuhan) (n=3) as reported.
**G.** Gating strategies for flow cytometry.

Figure S2

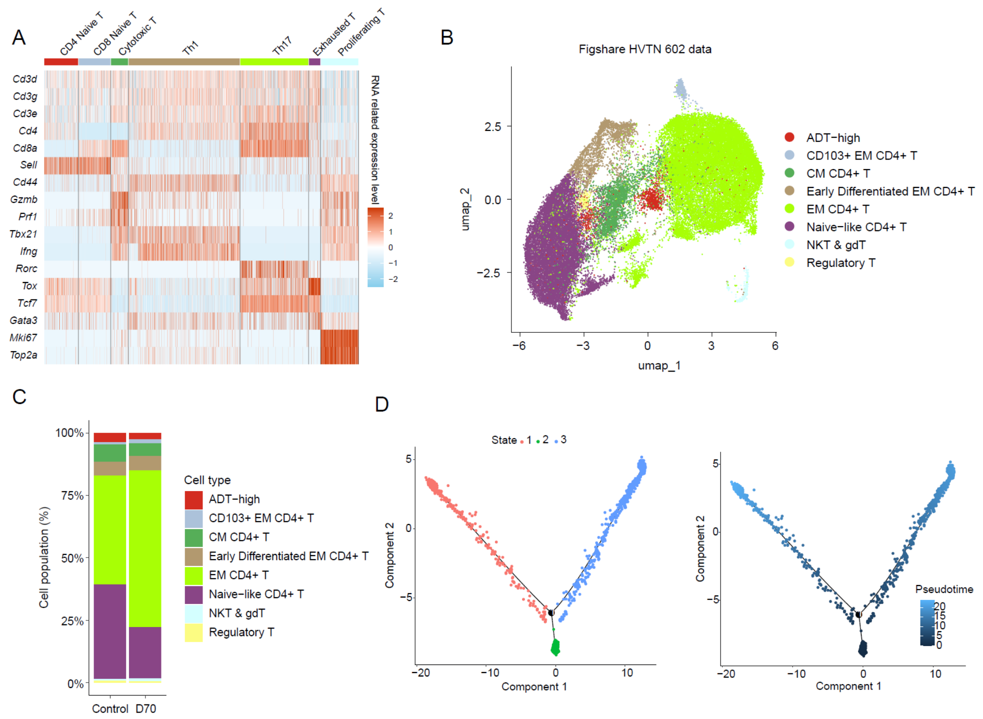


**Figure S2. T cell subtype identification and analysis of human BCG vaccination data from PRJNA1095450.**

**A.** Heatmap showing the expression levels of marker genes for different T cell subtypes.

**B.** Re-clustering of CD4$^+$ T cells from human BCG vaccination data, highlighting the clustering of T cell subtypes.

**C.** Bar plot showing the proportion of different CD4$^+$ T cell subtypes in the human BCG vaccination data.

**D.** Trajectory plot of the three T cell states, with distribution on the left and pseudotime progression direction on the right.

Figure S3

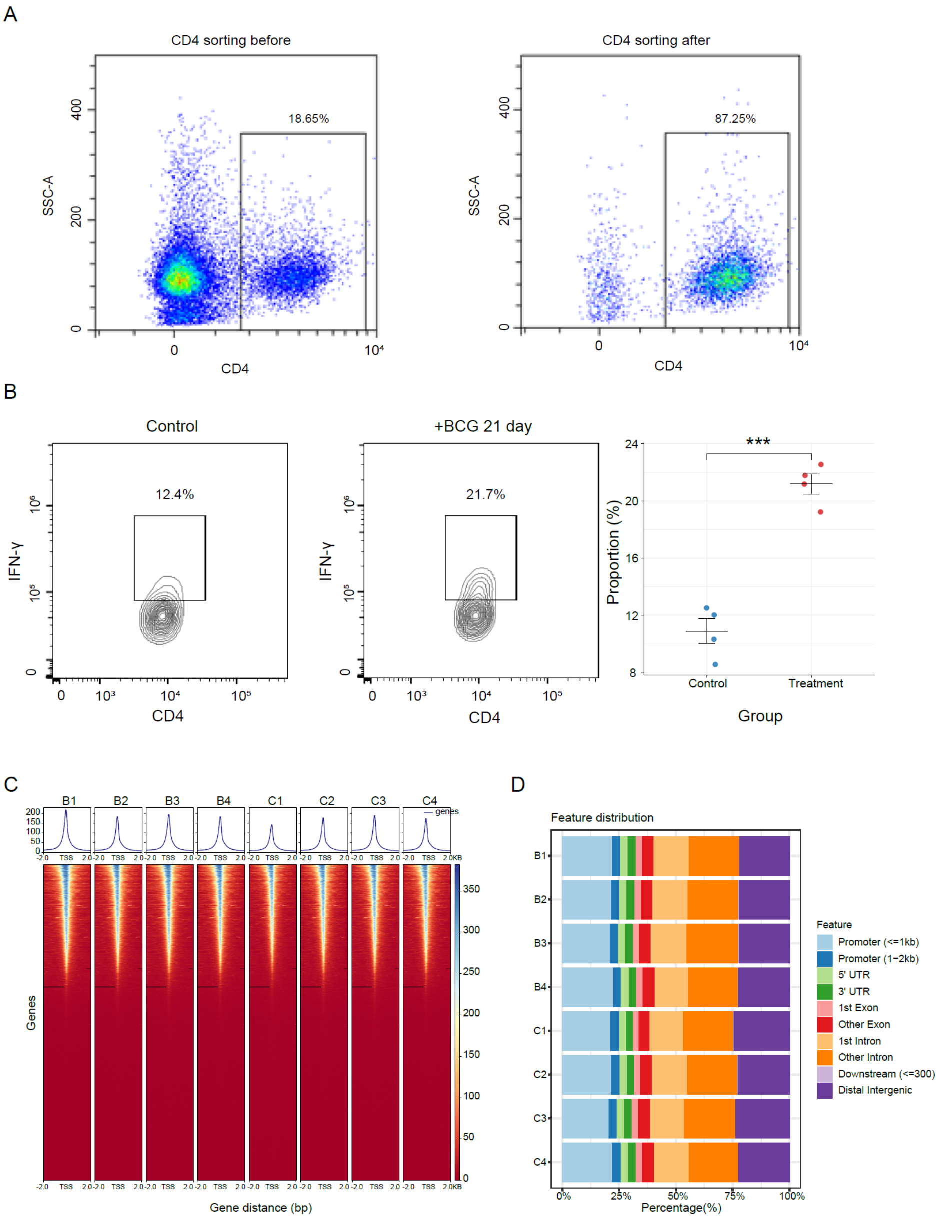
A
CD4 sorting before
18.65%
SSC-A
CD4
CD4 sorting after
87.25%
SSC-A
CD4
B
Control
12.4%
IFN-γ
CD4
+BCG 21 day
21.7%
IFN-γ
CD4
***
Proportion (%)
Control
Treatment
Group
C
B1
B2
B3
B4
C1
C2
C3
C4
genes
Genes
Gene distance (bp)
D
Feature distribution
Feature
Promoter (<=1kb)
Promoter (1−2kb)
5' UTR
3' UTR
1st Exon
Other Exon
1st Intron
Other Intron
Downstream (<=300)
Distal Intergenic
Percentage(%)

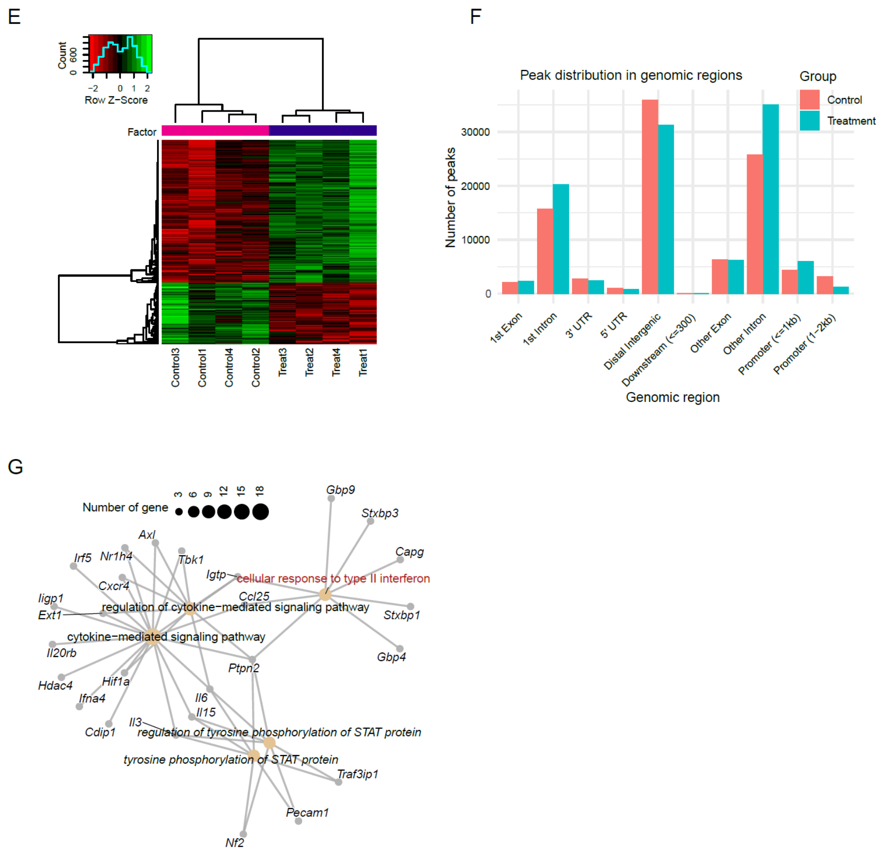


**Figure S3. ATAC-seq sample quality analysis and pathway enrichment display.**
**A.** Sorting efficiency of CD4$^+$ T cells as assessed by flow cytometry.
**B.** Flow cytometry analysis of IFN-γ$^+$ CD4$^+$ cells in the lungs of four groups of mice: control, 21 days post BCG, 21 days post BCG followed by 4 days of RSV infection, and 4 days of RSV infection only. *** $p < 0.001$.
**C.** Distribution of peaks within a region of ±2 kb around the TSS region (TSS4K) across the D21 and control groups. The plot demonstrates the consistency of peak distribution across the two groups within the gene promoter region.
**D.** Bar plot showing the distribution of peaks annotated to different genomic elements for each sample after peak calling. This analysis reveals how the peaks are distributed across various regulatory regions.
**E.** Comparison of binding sites between the D21 and control groups, showing the differences in chromatin accessibility between the two conditions.
**F.** Differential analysis of peaks with FDR < 0.05, indicating significant differences between the two groups. The analysis identifies the regions where the peaks are enriched after BCG immunization, shedding light on the chromatin regions most affected by the treatment.
**G.** The top 5 genes in the type II interferon signaling pathway, ordered by *p*-value. Genes related to the type II interferon response are highlighted, with the pathway marked in red.

Figure S4

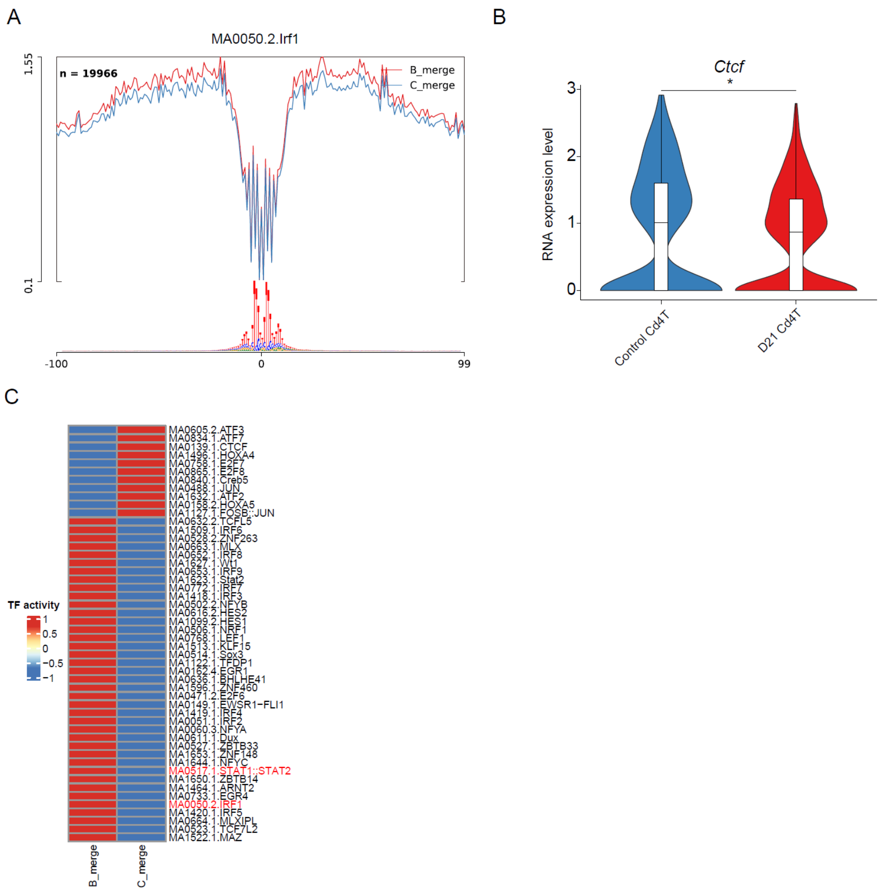


**Figure S4. TF footprint analysis across different sample groups.**

**A.** TF activity of Irf1 displayed after merging samples from both groups, highlighting its differential activity in the D21 and control groups.

**B.** Expression levels of *Ctcf* in CD4$^{+}$ T cells, displayed as a vlnplot, showing the variation in expression across different samples. * $p < 0.05$.

**C.** TF footprint analysis after merging different samples from both groups. The heatmap shows the TF activity of the top 50 TFs with the highest variance between the two groups.

Figure S5

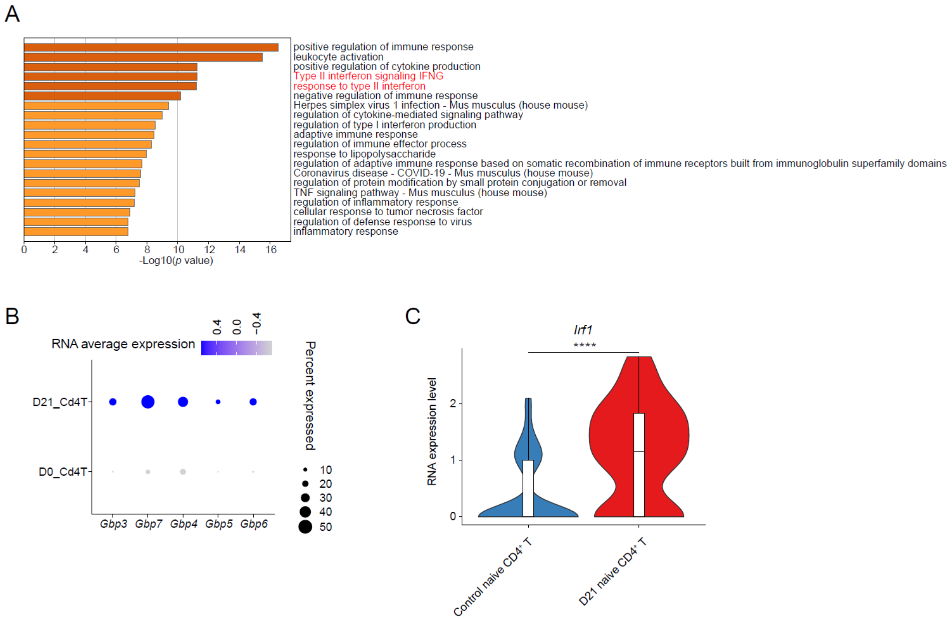


**Figure S5. Analysis of target genes regulated by the Stat1 regulon in CD4⁺ T cells**

**A.** Pathway enrichment analysis of genes regulated by Stat1, highlighting key pathways that these genes are involved in, such as those related to immune responses and interferon signaling.

**B.** Dot plot showing the expression of the Gbp family genes, which are regulated by Stat1, in CD4⁺ T cells. This includes genes like *Gbp3*, *Gbp4*, *Gbp5*, *Gbp6*, and *Gbp7*, which are upregulated following BCG immunization.

**C.** The violin plot of *Irf1* gene expression levels in naïve CD4⁺ T cell. **** $p < 0.0001$.

**SI table 1. Supplementary information on key resources***

| REAGENY or RESOURCE | SOURCE | IDENTIFIER |
|---|---|---|
| Antibodies | | |
| PE/Cyanine7 anti-mouse IFN-γ Antibody | BioLegend | Cat# 505826 |
| FIXABLE VIABILITY DYE EF780 100T | eBioscience | Cat# 65-0865-14 |
| CD45 Monoclonal Antibody (30-F11), eFluor™ 450 | eBioscience | Cat# 48-0451-82 |
| CD3e Monoclonal Antibody (145-2C11), APC | eBioscience | Cat# 17-0031-82 |
| CD4 Monoclonal Antibody (GK1.5), FITC | eBioscience | Cat# 11-0041-82 |
| Reagents | | |
| Brefeldin A Solution (1,000X) | BioLegend | Cat# 420601 |
| PMA/TPA | Beyotime | Cat# S1819 |
| Ionomycin | Beyotime | Cat# S1672 |
| Liberase™ TM Research Grade | Roche | Cat# 79439800 |
| DNaseI | Roche | Cat# 80218600 |
| Mouse IL-18 ELISA Kit | CUSABIO | CSB-E04609m |
| Mouse IFN-γ ELISA Kit | Proteintech | KE10094 |
| Mouse TNF-α ELISA Kit | CUSABIO | CSB-E04741m |
| Mouse $Cd4^{+}$ T Cell Isolation Kit | Vazyme | CS102 |
| Hyperactive ATAC-Seq Library Prep Kit for Illumina | Vazyme | TD711 |
| Virus and BCG | | |
| Human RSV type A | ATCC | Cat# VR-1540 |
| BCG live vaccine | Ruichu BIO | Lot: 20240718 |
| Experimental models | | |
| Mouse: BALB/c | SPF Biotech | Female mice (6 weeks of age) |
| Software and Algorithms | | |
| fastp | Chen | https://github.com/OpenGene/fastp |
| Sambamba | Tarasov, et al. | https://github.com/biod/sambamba |
| SAMtools | Danecek, et al. | https://github.com/samtools/samtools |
| BAMscale | United States Government Work | https://github.com/ncbi/BAMscale |
| MACS2 | Zhang, et al. | https://pypi.org/project/MACS2 |
| HOMER | Heinz, et al. | http://homer.ucsd.edu/homer/motif/ |
| HINT-ATAC | Li, et al. | https://reg-gen.readthedocs.io/en/latest/hint/introduction.html |
| DeepTools | Ramírez, et al. | https://test-argparse- |

| | | |
|---|---|---|
| | | readoc.readthedocs.io/en/latest/ |
| DiffBind | Stark, et al. | https://github.com/nshanian/DiffBind |
| ChIPseeker | Yu, et al. | https://github.com/YuLab-SMU/ChIPseeker |
| clusterProfiler | Wu, et al. | https://github.com/YuLab-SMU/clusterProfiler |
| Seurat | Hao, et al. | https://satijalab.org/seurat/ |
| CellChat | Jin, et al. | https://github.com/sqjin/CellChat |
| msigdbr | Dolgalev | https://github.com/igordot/msigdbr |
| SCENIC | Aibar, et al. | https://github.com/aertslab/SCENIC |
| GSVA | Hänzelmann, et al. | https://github.com/rcastelo/GSVA |
| Metascape | Zhou, et al. | https://github.com/data2code/Metascape |

* The sources and catalog numbers of key reagents are listed. The table also lists the sources and developers of software and algorithms.
